## Supplementary Figures for "Indistinguishable mitochondrial phenotypes after exposure of healthy myoblasts to myalgic encephalomyelitis or control serum"

### Supplementary Figure 1

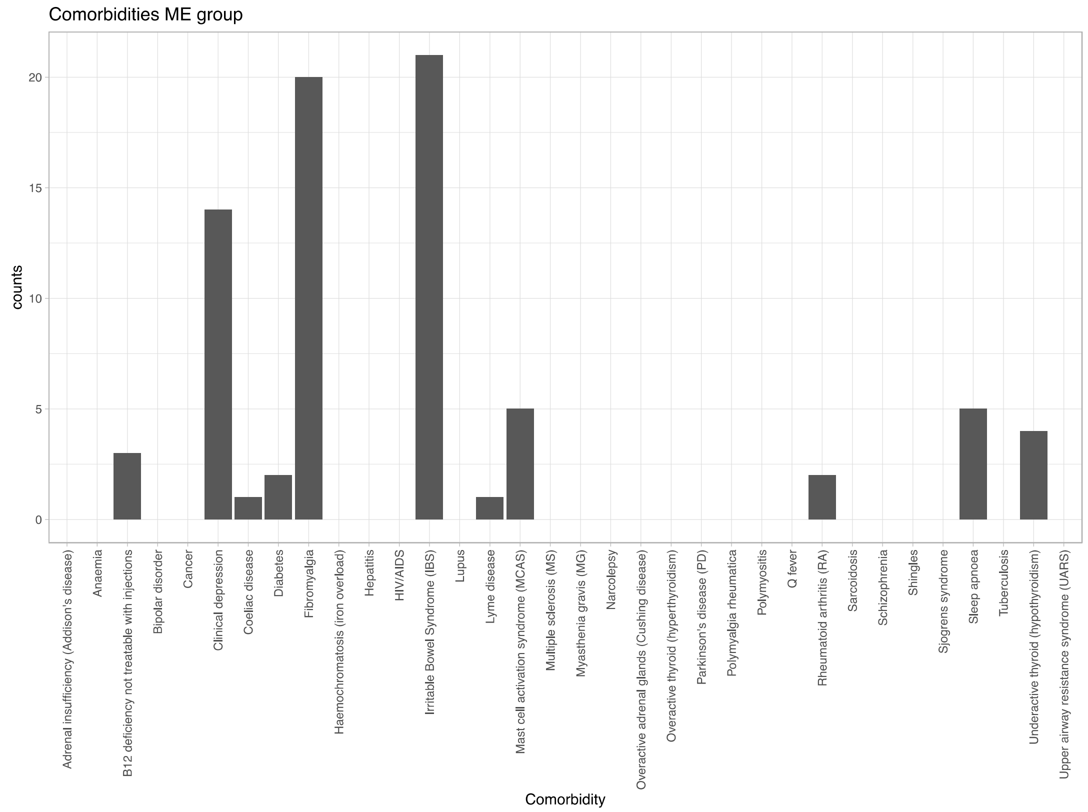


### Supplementary Figure 2: Testing oligomycin on cells treated with FBS only

3uM Oligomycin, 2 uM FCCP

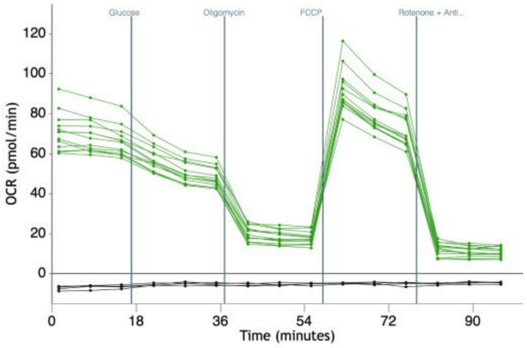


2uM Oligomycin, 2 uM FCCP

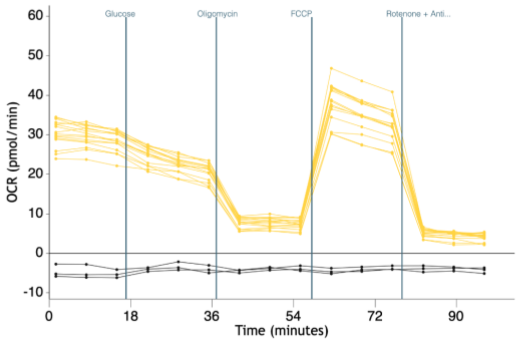
